# Cactaceae: uses and knowledge of rural communities in the semi-arid zones of Cucaita, Ráquira and Villa de Leyva (Boyacá, Colombia)

**DOI:** 10.1101/2020.12.08.416065

**Authors:** Daniela Porras–Flórez, Sofía Albesiano, Kendry Hernández–Herrera, Nubia Gómez–Velázco, Leopoldo Arrieta–Violet

## Abstract

Taxa of the family Cactaceae presents morphological and reproductive characteristics, which are used by rural communities in arid and semi-arid areas, for food, medicinal and ornamental purposes. The objectives were to identify the most used species and to relate the socioeconomic factors of the informants with their knowledge of the species. To this end, 262 semi-structured interviews were applied, with questions such as name, age, gender, educational level, source of employment, among others; eight categories of use were established: agro-ecological, agricultural, commercial, environmental service, food, medicinal, ornamental and technological; three indexes were calculated, relative importance, the value of use for each of the species and wealth of knowledge of the interviewees. Nine species are reported, from which eight are used as food and ornamental, standing out *Opuntia ficus-indica* for its diversity of uses, while *Cylindropuntia tunicata* does not report any utility. Variables such as age and residence time showed a significant relationship with the number of plants used by the interviewees. It is expected to contribute to the preservation of cacti, local knowledge and encourage large-scale cultivation since species such as *Mammillaria columbiana, Melocactus andinus*, and *Melocactus curvispinus* are being used in an unsustainable way, which could cause their local extinction.

## Introduction

Climate change, erosive processes, and anthropogenic activities such as agriculture, lumbering, and grazing have increased arid and semi-arid areas in Colombia, these areas have 4,828,875 hectares, which corresponds to 4.1% of the national territory, extending mainly along with the western, central, and eastern mountain ranges, and on the Caribbean coast, from sea level to 2,800 meters above sea level [1,2].

The department of Boyacá is affected by a serious desertification process with low sustainability, presenting semi-arid enclaves in the Central and Ricaurte provinces, in the municipalities of Cucaita, Ráquira, and Villa de Leyva, characterized by superficial soils, with highly eroded sectors, slopes of up to 60° and water deficit [3–5], conditions that prevent them from being productive areas, directly affecting the peasants’ economy. However, the largest extension of cultivated land corresponds to the pea *Pisum sativum*-Fabaceae, barley *Hordeum vulgare*, bulb onion *Allium cepa*, branch onion *Allium fistulosum*-Alliaceae, maize *Zea mays*-Poaceae, potato *Solanum tuberosum*-Solanaceae, and carrot *Daucus carota*-Apiaceae [6,7], which require constant humidity for high profitability, benefiting only farmers in humid areas.

The cacti of the *Austrocylindropuntia* and *Opuntia* generates stand out in the plant communities of the rural region of Cucaita, Ráquira, and Villa de Leyva, which present characteristics own for its development and propagation, such as their asexual reproduction, low water, and nutritional requirements for their growth, allowing them to occupy large extensions of land, and whose cultivation could become a source of income for the inhabitants of these municipalities, by using the native flora as an alternative for a more stable and lasting peasant economy, resulting in a higher index of food autonomy in these sectors.

Studies on knowledge and use of cacti in Colombia are limited; the food, medicinal and technological use of the leaves, fruits, and stems of *Acanthocereus pentagonus, Hylocereus undatus, Melocactus coccineus, Melocactus curvispinus, Opuntia schumannii, Pereskia bleo*, and *Stenocereus griseus* has been reported [8–11].

In municipalities with economic problems that lead to hunger and poverty, an understanding of the human-nature relationship is fundamental to the design of culturally appropriate and sustainable economic promotion and development projects [12]. Therefore, the objectives of this study were to identify the native species and among them the most exploited ones, to assign the categories of use, and to analyze the relationship between the socioeconomic factors of the interviewees and the knowledge of the taxa.

## Material and Methods

### Area of study

Three field trips were conducted, with a duration of five days each, between August and September 2019, to the semi-arid trails of the municipalities of Cucaita: Centro, Cuesta en Medio and Llano (5°31’55’’N, 73°27’44’’W; 2650 masl); Ráquira: Candelaria Occidente, Candelaria Oriente, Carapacho (5°31’12.22”N, 73°36’20.92”W; 2165 masl) and Villa de Leyva: Cañuela, Llano Blanco, Moniquirá, Ritoque, Sabana and Sopotá (5°36’16.19’’N, 73°31’34.92’’W; 2149 masl), in the department of Boyacá (Fig 1).

**Fig 1.**
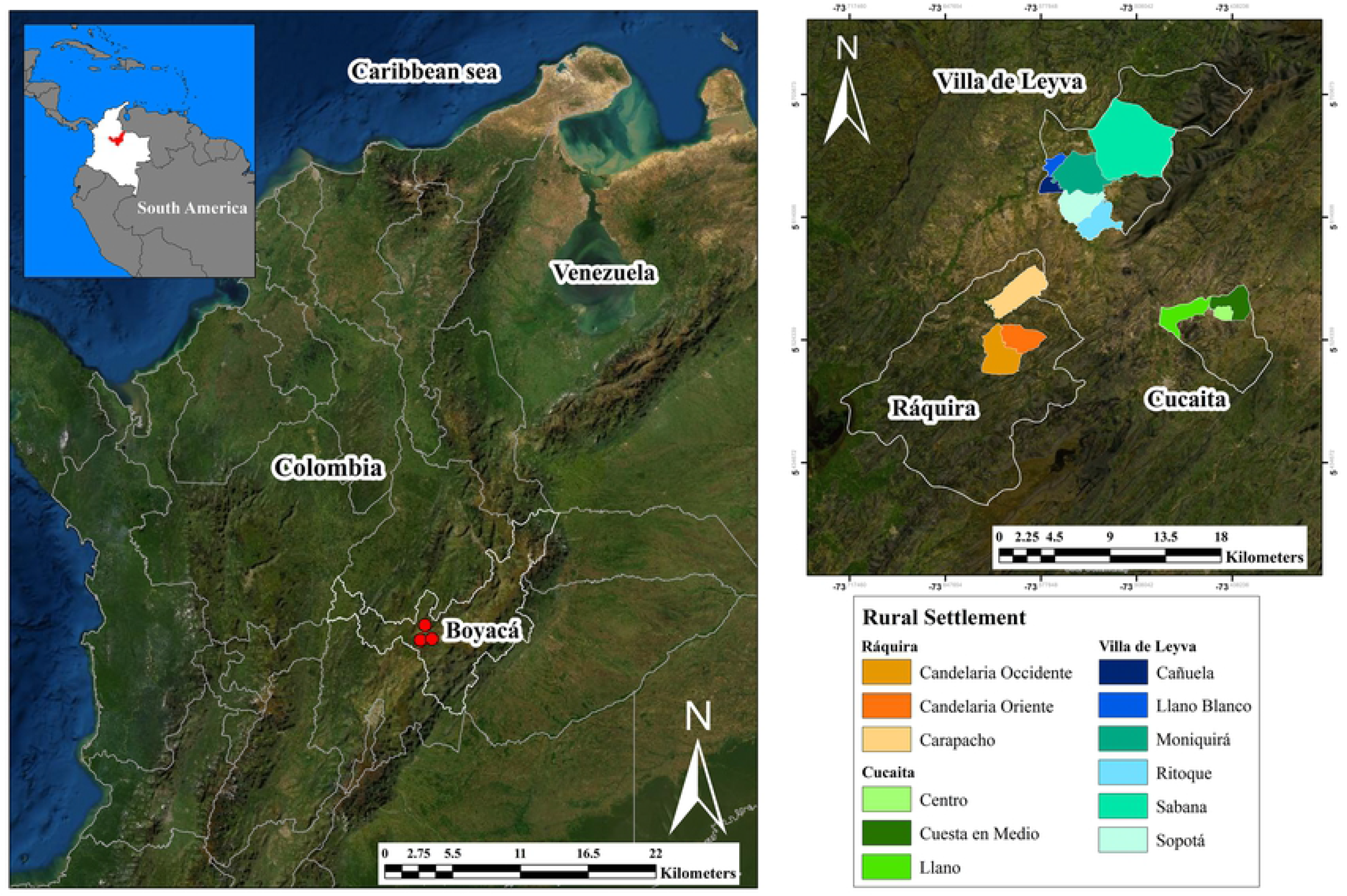
Study area. a. Municipalities located in the semi-arid inter-Andean enclaves of the Eastern Cordillera. b. Trails sampled in each municipality.

The three municipalities form a semi-Arid-Andean corridor, with temperatures between 14 and 18°C and an average annual rainfall of 700 to 1000 mm. The soils are clayey and sandy, and there are sectors of high erosion, with slopes of up to 60° and varied topography, between undulating and gullies [5,13–15].

The predominant type of vegetation corresponds to xerophytic and sub-xerophytic scrub, with *Agave americana* and *Furcraea* sp. -Agavaceae, *Puya nitida*-Bromeliacee, *Echeveria bicolor*-Crassulaceae, *Caesalpinia spinosa*-Caesalpiniaceae, *Xylosma spiculifera*-Flacourtiaceae, *Dodonaea viscosa*-Sapindaceae, *Solanum crinitipes, Solanum lycioides*-Solanaceae, and cacti of the genera *Mammillaria, Melocactus* and *Opuntia* [16–20].

### Specimen Collection

Specimens of the nine taxa were randomly collected and deposited in the UPTC Colombia herbarium of the Universidad Pedagógica y Tecnológica de Colombia, an institution that holds the Permit for the Collection of wild species of Biological Diversity the purpose of non-commercial scientific research, Resolution No. 724 of 2014, of the Autoridad Nacional de Licencias Ambientales–ANLA.

### Taxonomic identification

It was carried out based on the consultation of the taxonomic treatments of [19–21]. A photographic catalog was designed with the cactus species of each municipality, helping to facilitate the recognition of the taxa by the informants (S1 Appendix).

### Ethical aspects

This study was approved according to the guidelines and regulations of the Ethics Committee for Scientific Research of the Universidad Pedagógica y Tecnológica de Colombia (Agreement 063 of 2016). Each informant declared to know the objectives and methods of the project, accepting their voluntary participation by signing the Informed Consent Form, allowing the publication of ethnobotanical information (S2 Appendix).

### Estimation of the sample size (n) people to be interviewed

The number of inhabitants per village (N) was calculated according to the number of rural inhabitants over 18 years old in each municipality and the population density [22]. The formula from [23] was applied to determine the statistically representative number of people to be interviewed (n) per municipality:

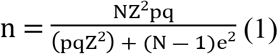

where, n= sample size. N= estimated number of inhabitants of the semi-arid trails. Z= 1.96, confidence level = 95%, a probability of success of p= 0.5, which it involves q= 0.5, permissible error, e= 0.1.

Given the difficulty of having a sampling frame, the non-probabilistic type of sampling was applied under a criterion of proximity and by quotas [24], where the people who reported living in the study area were selected.

### Structure and application of the interview

A semi-structured interview was used, with two groups of questions, personal data, and ethnobotanical information: name, age, gender, educational level, source of employment, time spent in the area, common or vernacular name of each species, uses, manner of employment and structure of the plant used (S2 Appendix), each person was assigned an identification code, and the respective depuration and normalization was carried out to form the database.

The results of the interview were analyzed according to the categories proposed by [25], defined as:

**Agro-ecological:** they provide benefits to agro-ecosystems, such as living fences, compost, soil protection, among others. **Agricultural:** they fulfill an agro-industrial function to facilitate agricultural and/or livestock processes, such as fodder, sowing tools, and veterinary. **Food:** they provide food to the inhabitants. **Commercial:** they are placed exclusively for sale. **Medicinal:** they have preventive and/or curative properties for human diseases. **Ornamental:** due to their striking phenotype they are used to decorate the interior and/or exterior of houses. **Environmental services:** they are wild and according to the local inhabitants they are associated with the recovery of ecosystems or the preservation of fauna, for example, reforestation and food for wild animals. **Technological:** when transformed they provide mechanical or chemical aids for the daily activities of human communities; they can generate innovative products.

### Ethnobotanical Indices

Three indices were estimated: Wealth of Knowledge (RQZi) [11], Relative Importance (RI), and Use Value (UVs) [26]:

### Rich Knowledge Index (RQZi)

is the knowledge that the informant (i) has about the species in relation to all the species found in each municipality.

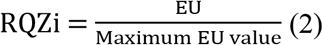

where, EU= is the total number of species recognized by the informants. Maximum EU value= Total number of species present in the area.

The RQZi index varies between 0 and 1, with 0 being the minimum, that is, they do not recognize any plant and 1, the maximum value of knowledge, know all the species present in the territory.

### Use Value Index (UVs)

measures the socio-cultural importance that the species has for an informant.

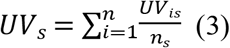

where, UV_is_= used value of species *(s)* to informant = *i*.

UV_is_= ∑U_*is*_/n_*is*_, where, U_*is*_= number of uses of species *(s)* mentioned in each event by informant *i*; n_*is*_= number of events in which informant *i* cited species. *s*n_s_= number of informants interviewed for species. *i*: informant. *s*: species.

### Relative Importance Index (RI)

reflects the utility of the plant according to the versatility of uses; the maximum value that can be assigned to a species is 2.

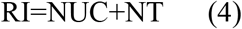

where, NUC= number of categories of use of the species/number of categories of use of the most used species

NT= number of uses attributed to the species/number of uses attributed to the most used species.

### Relationship of ethnobotanical data and socioeconomic factors

The study carried out is of a descriptive correlational type, after standardization, depuration, and codification of the information. Exploratory analyses were carried out in order to identify potentially atypical values through the Box Plot diagram. Univariate and bivariate analysis techniques were applied according to the number of plants used per municipality, part of the plant used, category of use and knowledge wealth index, variables that were crossed to determine the relationship or association with respect to a municipality, gender, occupation, and age of the interviewee.

The association between categorical qualitative variables, municipality, gender, and occupation with quantitative variables was determined through the Chi-square test of independence. Being the test statistic:

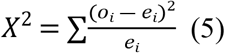

where, o_i_= observed frequencies. e_i_= expected frequencies. The degrees of freedom are determined by (f-1) (c-1), where f= number of rows in the contingency table, and c= number of columns in the contingency table.

For associations where both variables are quantitative, the Pearson Correlation Coefficient (R_x,y_) was calculated, after reviewing the required assumptions, including normality.

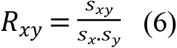

where, S_xy_= covariance between variables x, y. S_x_= standard deviation of x. S_y_= standard deviation of y.

Statistical hypothesis: null hypothesis. H_0_: no relationship between variables. Alternate hypothesis. H_1_: if there is a relationship between the variables.

For the hypotheses of no relation or association between the variables, both for the Chi-square statistic and for the R_xy_ Correlation Coefficient, the P value is calculated, considering three reference points for comparison: α =10% (*), α =5% (**), α =1% (***).

## Results and Discussion

### Cactus species present in the study area

Nine species were found (Table 1, Fig 2), which had been reported for the municipalities of Ráquira and Villa de Leyva by [5,11,18–20,27]; however, they were not observed in the sampling area *Disocactus* sp, *Melocact*us *hernandezii, Opuntia schumannii*, and *Rhipsalis baccifera*, described for Villa de Leyva and its surroundings by [5,11].

**Table 1.** Vernacular names and number of citations of species in the area.

| Species | Vernacle | Collection Number/Herbarium Number | Citations Number |
| --- | --- | --- | --- |
| **Cactoideae** |  |  |  |
| *M. columbiana* | Ajicitos, Cauto, Piña | KH4/UPTC-H 30070 | 299 |
| *M. andinus* | Ajicitos, Cauto | FV4/UPTC-H 30073 | 114 |
| *M. curvispinus* | Ajicitos, Cauto | FV6/UPTC-H 30074 | 114 |
| **Opuntioideae** |  |  |  |
| *A. cylindrica* | Penco, Cactus, Cacho de Buey | KH3/UPTC-H 30067 | 104 |
| *A. subulata* | Penco, Cactus, Cacho de Buey | DP98/UPTC-H 30068 | 104 |
| *C. tunicata* | Vanilla | Dp101/UPTC-H 30069 | - |
| *O. ficus-indica* | Higo, Higo de castilla | KH6/UPTC-H 30071 | 341 |
| *O. quitensis* | Higo silvestre, Higo, Penco de monte | DP7/UPTC-H 30072 | 70 |
| ***O. soederstromiana*** | Higo silvestre, Higo, Penco de monte | DP95/UPTC-H 30066 | 244 |

In the subfamily Cactoideae, *M. columbiana*, records the highest number of citations; in the subfamily Opuntioideae, *O. ficus-indica*. The least cited species was *O. quitensis*. KH: Kendry Hernández, DP: Daniela Porras, FV: Felipe Vargas. (-) Not applicable.

**Fig 2.**
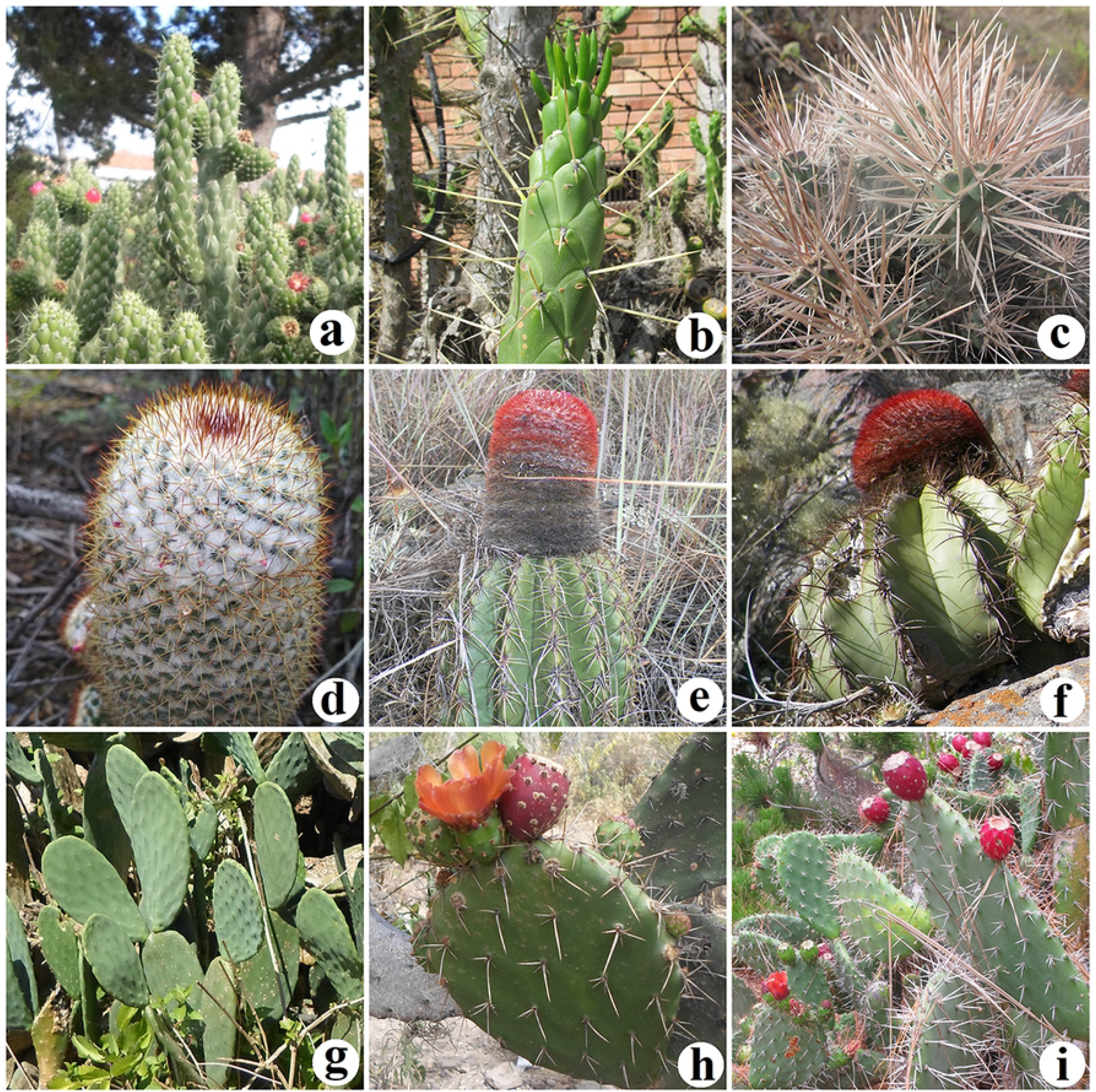
Species reported for the study area. a. *A. cylindrica*. b. *A. subulata*. c. *C. tunicata*. d. *M. columbiana* e. *M. andinus*. f. *M. curvispinus*. g. *O. ficus-indica*. h. *O. quitensis*. i. *O. soederstromiana*. Photographer: Daniela Porras-Flórez, a-d, g-i; Felipe Vargas: e, f.

### Uses attributed to cacti

The most reported use category was as food (661 citations), followed by ornamental (251 citations) and medicinal (138 citations). These categories are frequently mentioned in various studies in America, for example, in Mexico, species such as *O. ficus-indica* were fundamental to the establishment of large cultures such as the Aztec, since it served them mainly as food, and later was given other uses: medicinal, ornamental and forage for animals [28,29]. Currently, the constant consumption and commercialization of its fruits and stems have contributed to the internal economic development of that country [30].

Also, in different latitudes of the Americas, cacti have been part of the daily and traditional activities of communities, for example, indigenous peoples in Canada consume the fruits of *Escobaria vivipara* and the stems of *Opuntia fragilis* as food and to treat skin infections. In the United States, indigenous communities used *Opuntia humifusa* for medicinal purposes, to treat wounds produced by snake bites; today, this species is used as an ornamental [31]. In Cuba, Honduras, and Guatemala, species of the genera *Brasiliopuntia, Disocactus, Epiphyllum, Hatiora, Hylocereus, Mammillaria, Melocactus, Opuntia, Pereskia, Rhipsalis, Selenicereus, Stenocereus*, are frequently reported as ornamental. In Bolivia the genera *Cleistocactus, Disocactus, Epiphyllum, Pfeifera*, and *Rhipsalis*, and in Brazil species such as *Cereus jamacaru, Pilosocereus pachycladus, Pilosocereus gounellei*, are mainly used as livestock feed [30,32–40].

### Cactoideae subfamily

*M. columbiana* is known by its vernacular name of piña, due to the appearance of the ribs on the stem; the fruits are known as ajicitos and in Mexico as chilitos, because of their color and shape [28] (Table 1). 53% of the informants consume the fruits occasionally (136 citations), for their sweet flavor and striking color, which coincides with Mexico [28] (Table 2). In Villa de Leyva and Sáchica, the populations of this species have been decimated by the strong fires that occur periodically in the upper parts of the mountains of these municipalities, and by commercialization for ornamental use [41]. *M. columbiana* is distributed from Central America to northern South America; in Colombia, it is found in the inter-Andean valleys of the departments of Boyacá, Cundinamarca, Huila, and Santander [20,42,43].

**Table 2.** Categories and distribution of species.

| Species | Use category | Use | Utilization | Used part | C | R | V |
| --- | --- | --- | --- | --- | --- | --- | --- |
| **Cactoideae** |  |  |  |  |  |  |  |
| *M. columbiana* | Co | Sold in pot | Sold in Pot | CP | x |  | x |
|  | Fo | Sporadic consumption | In nature | FR | x | x | x |
|  | Or | Decorative pots | - | CP | x | x | x |
| *M. andinus* | Fo | Sporadic consumption | In nature | FR | - | x | - |
|  | Or | Decorative pots | - | CP | - | x | - |
| *M. curvispinus* | Fo | Sporadic consumption | In nature | FR | - | x | - |
|  | Or | Decorative pots | - | CP | - | x | - |
| **Opuntioideae** |  |  |  |  |  |  |  |
| *A. cylindrica* | Agr | Living fence | - | CP | x |  | x |
|  | Me | Cough treatment | Infused Petals | FL | x |  |  |
|  | Or | Decorative pots | - | CP | x | x | x |
|  | Tec | Water purification | Mucilage is boiled with water | ST |  |  | x |
| *A. subulata* | Agr | Living fence | - | CP | x | x | x |
|  | Med | Cough treatment | Infused Petals | FL | x |  |  |
|  | Or | Decorative pots | - | CP | x |  | x |
| *O. ficus–indica* | Agt | Veterinary to treat cattle indigestion | Cladodes are liquefied | ST | x |  |  |
|  |  | Cattle fodder | In nature or the cladodes are cut into small pieces | ST | x |  |  |
|  | Agr | Living fence | - | CP | x | x | x |
|  |  | Composting | Dolomite lime or worms are added to the stems | ST | x |  |  |
|  | Co | Stems are sold to promote new crops | - | ST | x |  |  |
|  | EnS | Decrease in eroded soils | Sow | ST |  | x |  |
|  | Fo | Consumption of the fruit, desserts, salads, ice cream | In nature or based on fruit pulp | FR | x | x | x |
|  |  | Salads | Cooking | ST |  | x | x |
|  | Med | Cough treatment | The petals are infused | FL |  | x | x |
|  |  | Treating pulmonary conditions | In nature or in juice | FR | x |  | x |
|  |  | Treating the flu |  |  | x |  | x |
|  |  | Controlling blood pressure |  |  | x |  |  |
|  |  | Cleansing the colon |  |  | x |  |  |
|  |  | Treating arthritis |  |  |  | x |  |
|  |  | Managing gastritis |  |  |  | x | x |
|  |  | Muscle pain and deflation | Poultice | ST |  | x | x |
|  |  | Treating pulmonary conditions | Liquefied Mucilage |  | x | x | x |
|  |  | Treating the flu |  |  | x | x | x |
|  |  | Lose weight |  |  | x |  |  |
|  |  | Cancer prevention |  |  | x | x | x |
|  |  | Treating arthritis |  |  |  | x |  |
|  |  | Cleansing the colon |  |  |  | x |  |
|  |  | Fever management |  |  |  | x |  |
|  |  | Managing diabetes |  |  | x |  | x |
|  |  | Controlling blood pressure |  |  | x |  |  |
|  | Or | Decoration | - | CP |  | x | x |
|  | Tec | Mask for hair or as a base for shampoo | The stem is liquefied and mixed with other ingredients | ST | x | x |  |
|  |  | Water purification | Boils along with contaminated water |  |  |  | x |
| *O. quitensis* | Fo | Sporadic consumption | In nature | FR | - | x | - |
|  | Med | Treat cough | Infused Petals | FL | - | x | - |
| *O. soederstromiana* | Agt | Cattle fodder | - | CP | x |  | x |
|  | Agr | Living fence | - | CP | x |  | x |
|  | EnS | Wild bird food | In natura | FR | x |  | x |
|  | Fo | Sporadic consumption | In natura | FR | x | x | x |
|  | Med | Treating pulmonary conditions | Infused Petals | FL | x |  | x |
|  |  | Cleansing the colon | Liquefied Mucilage | ST | x | x | x |
|  |  | Treating varicose veins | Poultice |  |  |  | x |
|  |  | Back pain |  |  |  |  | x |

*Cylindropuntia tunicata* is not included because informants did not report uses. Use category: Agriculture (Agt), Agroecological (Agr), Commercial (Co), Environmental service (EnS), Food (Fo), Medicinal (Med), Ornamental (Or), Technological (Tec). Used part: Complete plant (CP). Flower (FL). Fruit (FR). Stem (ST). Not applicable (-). Presence in the municipality (x). Abbreviation: Cucaita (C), Ráquira (R), Villa de Leyva (V).

Informants record the same uses for *Melocactus andinus* and *M. curvispinus*, because of their morphological similarity they cannot differentiate them. In Ráquira, 34% (28 citations) of residents use them mainly as ornamentals. The Wayuú indigenous in the department of La Guajira, in addition to consuming the fruit of *M. curvispinus*, use the pulp as a source of water, as the base of dough to prepare arepas and to make syrup [44] and in Venezuela they also use it as food [45]. The specific epithet of *Melocactus andinus* refers to its distribution in the Andes of Colombia and Venezuela, while *M. curvispinus* refers to the curved shape of its spines [19,46]. In Colombia, *M. curvispinus* is known by several vernacular names, cabecinegro, cabeza de indio, cabeza de negro, and gorro de obispo [47].

### Opuntioideae subfamily

More than 50% of the interviewees consider *A. cylindrica* and *A. subulata* to belong to the same species, as they share a high number of morphological characters. Between 12 and 25% of the inhabitants mentioned ornamental use (57 citations), and between 3 and 6% as a living fence (9 and 7 citations respectively) (Fig 3a, Table 2), which coincides with what was registered for Tunja [48]. In Perú, they are cultivated as a living fence, and formerly the spines were used as needles for sewing and as comb [29,39]. The etymology of *A. subulata* refers to the shape of its deciduous leaves, while *A. cylindrica* refers to the shape of its stems. In Perú, *A. subulata* subspecies *exaltata* is known as “uchu uchu” and in Colombia, the inhabitants distinguish it as Cacho de Buey (Ox Horn) due to the curved shape of some of its branches (Table 1). These species are distributed throughout the Andean region in Bolivia, Colombia, Ecuador, and Peru, except *A. cylindrica* which is not present in Bolivia [43].

**Fig 3.**
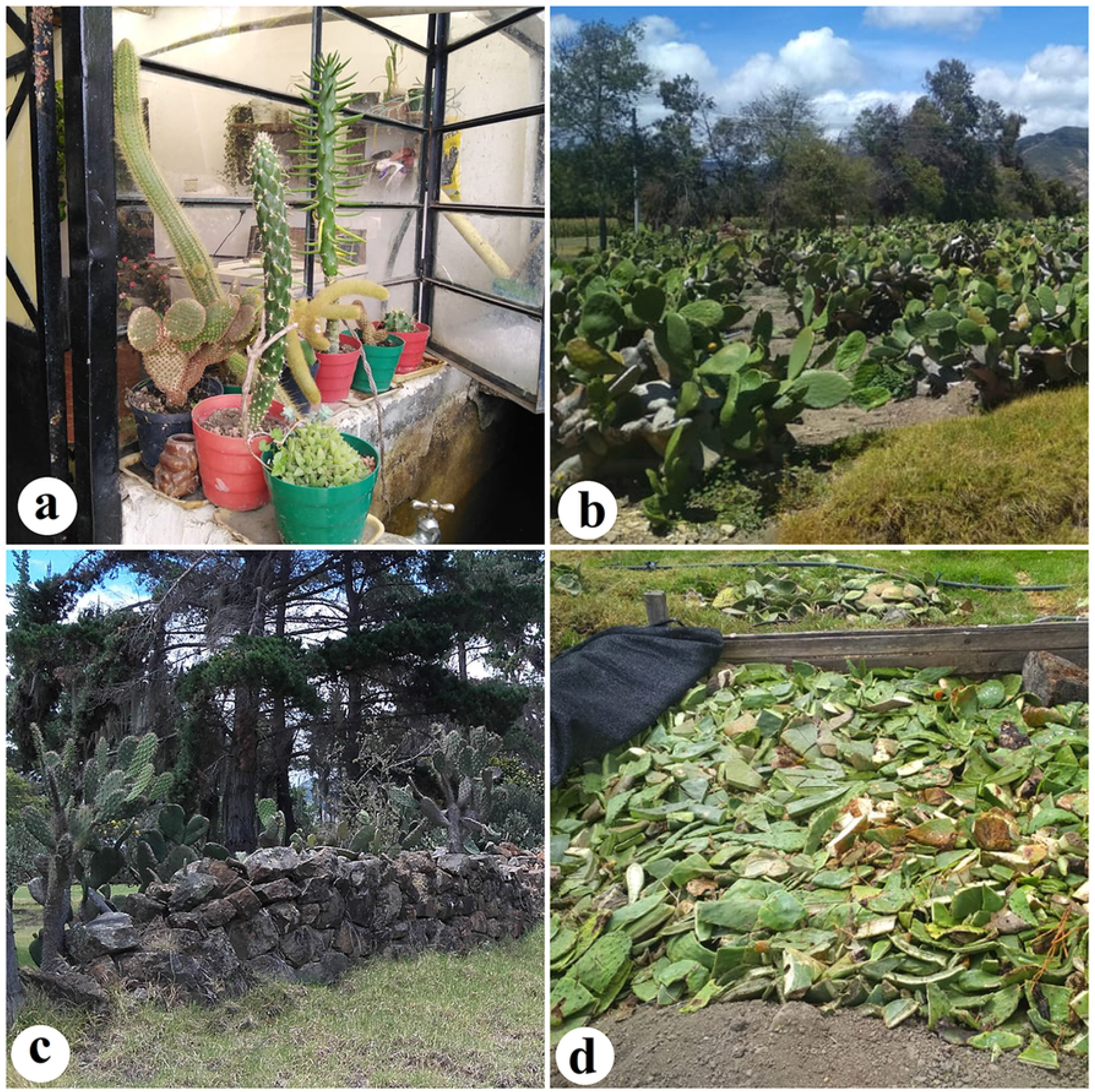
Some uses by the community are studied. a. Plantpot with young plants of *A. cylindrica* and *A. subulata* along with other introduced species. b. Cultivation of *O. ficus-indica* in the municipality of Cucaita (Boyacá). c. *O. soederstromiana* as a live fence. d. Cladodes of *O. ficus-indica* in decomposition for composting. Photographer: Daniela Porras-Flórez, a-d.

In Villa de Leyva *C. tunicata* it is known as Vanilla and no use is assigned to it, due to the dispersion of the branches by zoochory, when there is an increase in temperature, the spines become embedded in the skin of the animals and peasants, generating pain and discomfort [49]; therefore, the inhabitants have tried to eradicate it. The specific epithet of *C. tunicata* refers to the papery epidermal sheaths on the spines. This species has a wide Neotropical distribution, from the south of the United States to the north of Argentina [50].

Four crops of *O. ficus-indica* were found in the study area for the commercialization of the cladodes and fruits (Fig 3b). Besides, the discarded stems are used as organic fertilizer (composting) (Fig 3d), which shows that the species is being valued in the region, which coincides with the agricultural, agroecological, nutritional and medicinal uses that it has in other countries [29,35,51]. *O. ficus-indica* is used between 68 and 87% as a food (230 citations), especially the fruits in Cucaita, Ráquira, and Villa de Leyva.

The etymology of *O. ficus-indica* refers to its fruit, *ficus* corresponds to the higo, and *indica* to the West Indies [39]. In Colombia it is known as cactus, higo, penca de Castilla, and tuna [47], in the area of study is commonly called higo de Castilla, according to [52], this name is because Castilla was the first European center where the species was disseminated, in Bolivia as airampu or tuna, in Peru as the higo de las Indias. It is naturally distributed in the Neotropical region, from Canada to Patagonia in Argentina, and has been introduced for food, forage, and ornamental purposes in the Southern, Holarctic, and Paleotropical regions [53,54].

43% of the rural community interviewed (64 citations), sporadically consume the fruits of *O. quitensis*, and express that the presence of abundant glochids makes it difficult to manipulate them. Some farmers were able to distinguish it from the other *Opuntia* species because of its smaller fruits, others confused it with *O. soederstromiana* and therefore reported the same categories of use. In Ecuador, it is known in the province of Loja as Tuna and Tunilla, and its fruits are also consumed [55,56]. This species is distributed in Colombia, Ecuador, and Perú [21,57,58].

According to the inhabitants interviewed, *O. soederstromiana* is similar to *O. ficus-indica*, but with more spines, sweeter and smaller fruit, and they assign the same properties to it, therefore, they report some similar uses. 64% of people occasionally consume the fruit (185 citations) and 4% use it as a living fence (7 citations) (Fig 3c); coinciding with what has been registered in Ecuador, however, there they attribute other medicinal properties to it, since, according to popular knowledge, the roots serve to treat some stomach ailments and the mucilage of the cladodes is used as an expectorant, to treat fever and skin spots [59].

The specific epithet of *O. soederstromiana* is due to the first collector, Ludovic Söderstrom; it is found in Colombia [27] and is commonly known as higo silvestre and in Ecuador, it is known as tuna de San Antonio or pishku tuna.

## Ethnobotanical Indexes

### Knowledge wealth index (RQZi)

The knowledge of the species in the three municipalities is low, in Cucaita and Ráquira 16% and 10% respectively know all the species. Between 10% and 40% of the people reached an RQZi index of 0.3 to 0.6 (Fig 4).

**Fig 4.**
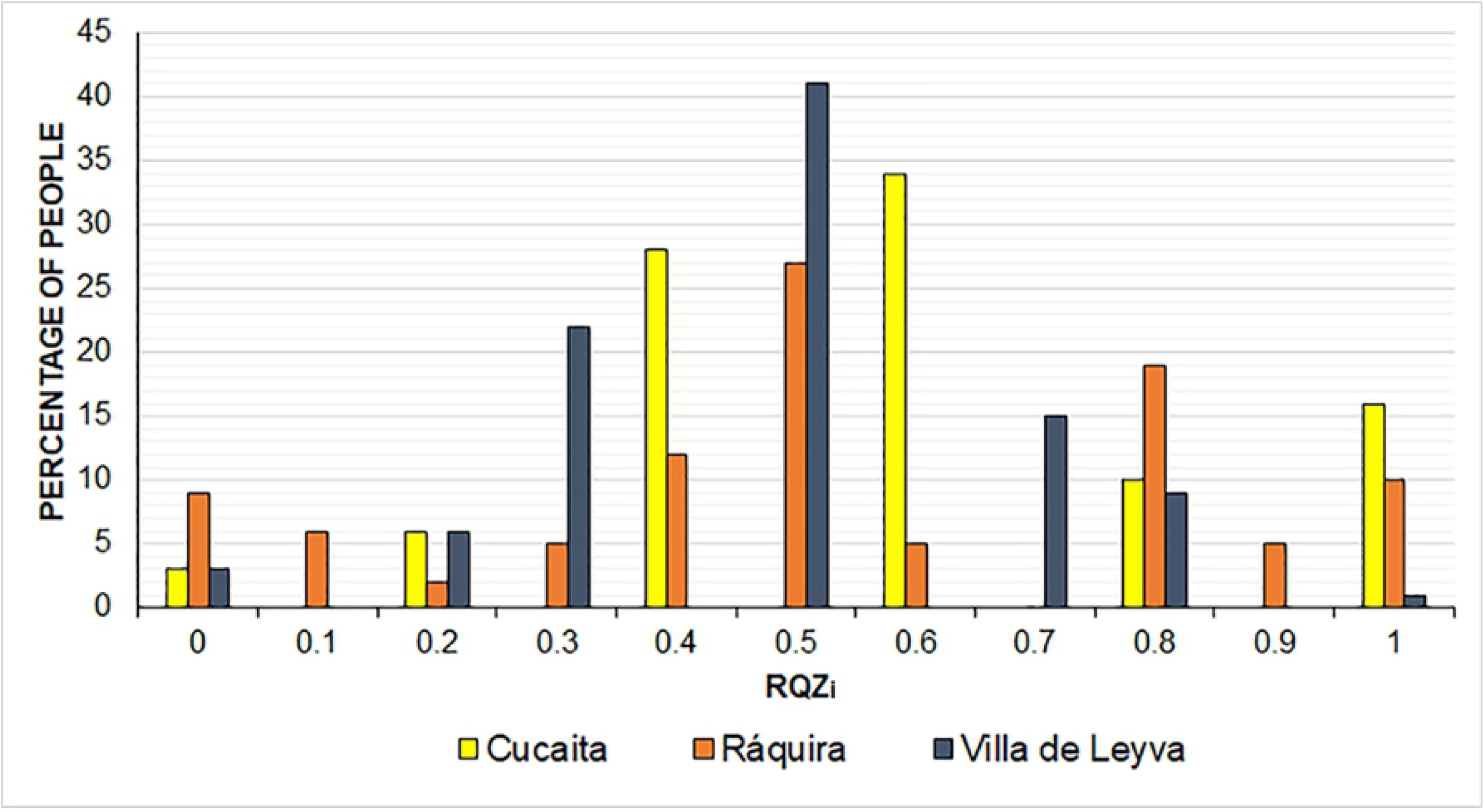
Informants knowledge wealth Index of by municipality. The maximum value corresponds to 1, which means the interviewee recognized all the species in his/her municipality.

### Relative importance index (RI)

*O. ficus-indica* had the highest value, 2, in the three municipalities (Table 3), because 28 uses were reported, grouped in the eight categories. This species is considered the most economically valuable in the world [52]. It is cultivated mainly for human consumption and medicine and as forage in Argentina, Bolivia, Brazil, Chile, Cuba, Guatemala, Honduras, Nicaragua, Dominican Republic, and for the production of the cochineal insect (*Dactilopius opuntiae*-Dactylopiidae), which is economically important in Ecuador and Peru for obtaining the main natural carmine red pigment [32,34,37,39,54,60–64].

**Table 3.** Use value and relative importance indexes for each municipality.

| Species | USE VALUE<br>(UVs) |  |  | RELATIVE IMPORTANCE<br>(RI) |  |  |
| --- | --- | --- | --- | --- | --- | --- |
|  | C | R | V | C | R | V |
| **Cactoideae** |  |  |  |  |  |  |
| *M. columbiana* | 0.9 | 0.74 | 0.82 | 0.7 | 0.41 | 0.87 |
| *M. andinus* | - | 0.63 | - | - | 0.41 | - |
| *M. curvispinus* | - | 0.63 | - | - | 0.41 | - |
| **Opuntioideae** |  |  |  |  |  |  |
| *A. cylindrica* | 0.27 | 0.26 | 0.2 | 0.97 | 0.2 | 0.87 |
| *A. subulata* | 0.26 | 0.26 | 0.3 | 0.7 | 0.2 | 0.58 |
| *C. tunicata* | - | - | 0 | - | - | 0 |
| *O. ficus-indica* | **1.85** | **1.06** | **1.06** | **2** | **2** | **2** |
| *O. quitensis* | - | 0.45 | - | - | 0.41 | - |
| *O. soederstromiana* | 0.94 | 0.86 | 1 | 1 | 0.41 | 1.7 |

The highlighted numbers correspond to the maximum value obtained among the species. (-) Not present in the municipality. Cucaita (C), Ráquira (R), and Villa de Leyva (V).

### Use Index Value (UVs)

In the three municipalities, *O. ficus-indica* registered the highest use value because it is the only species cultivated, and its cladodes and fruits are sold locally and nationally, indicating the species is becoming known better and used in rural communities. A value of 1.85 in Cucaita is the most outstanding (Table 3); its use at the local level can be influenced by its morphological characteristics, absence or few spines, cladodes, and larger fruits, besides being well positioned in international markets. *O. soederstromiana* was the second most used species, these results agree with [59] in Ecuador, where it is used for several purposes; this species has advantages that could introduce it in the local market, as its sweeter fruit and is also dominant in its ecosystem, and could be used as a natural barrier against winds causing erosion in the area.

### Ethnobotanical data and socioeconomic factors

Several studies that integrate ethnobotany and socioeconomic factors at the global level, not only for the conservation of biodiversity and local knowledge but also for the benefits in the quality of life that plants bring to communities [65–68].

Several ethnobotanical studies have focused on the influence of socioeconomic factors on knowledge and uses of natural resources [69] since it has been shown that there is a complexity of variables that influence human behavior concerning its management. Socioeconomic analysis of variables has increased the understanding of factors that affect the knowledge and use of native plants, which helps to identify better strategies to favor regional sustainability [67].

Correlations were made between the number of use categories mentioned, the number of plants used, and the parts used, age, gender, level of studies, occupation, and residence time living in the area (Table 4).

**Table 4.**
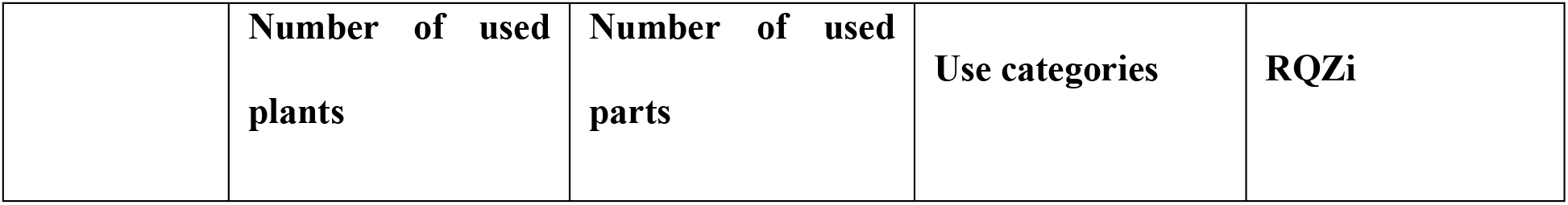
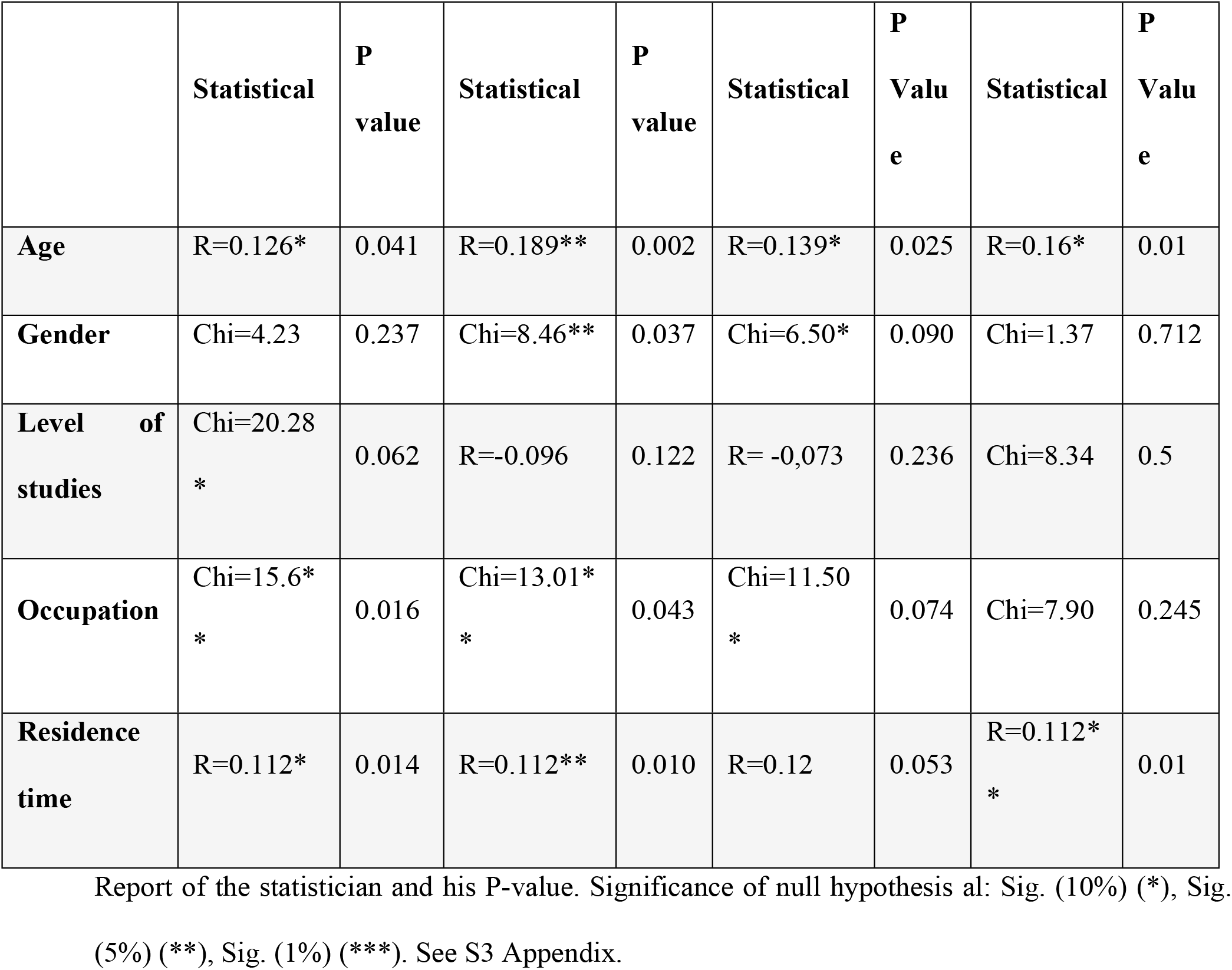
Summary of relationships between ethnobotanical data and socioeconomic factors.

### Age

Pearson’s correlation coefficient shows that there is a statistically significant correlation at 5%, between the age of the interviewee and the knowledge index (R=0.16), as well as with the categories of use, the number of plants used and the number of parts used (Table 4). There is evidence of a slightly higher level of knowledge among respondents aged 50 or more, and a lower level of knowledge among those under 30 (Fig 5). These results coincide with those reported by [70], who maintain that people between the ages of 55 and 90 are the greatest holders of local knowledge, due to the long history of contact that has favored the exchange of knowledge and experiences created with plants throughout their lives [71]. [72] point out that access to technologies by young people such as television, cell phones, radio, among others, has decreased their interest in outdoor activities, therefore their relationship with local biodiversity is weak.

**Fig 5.**
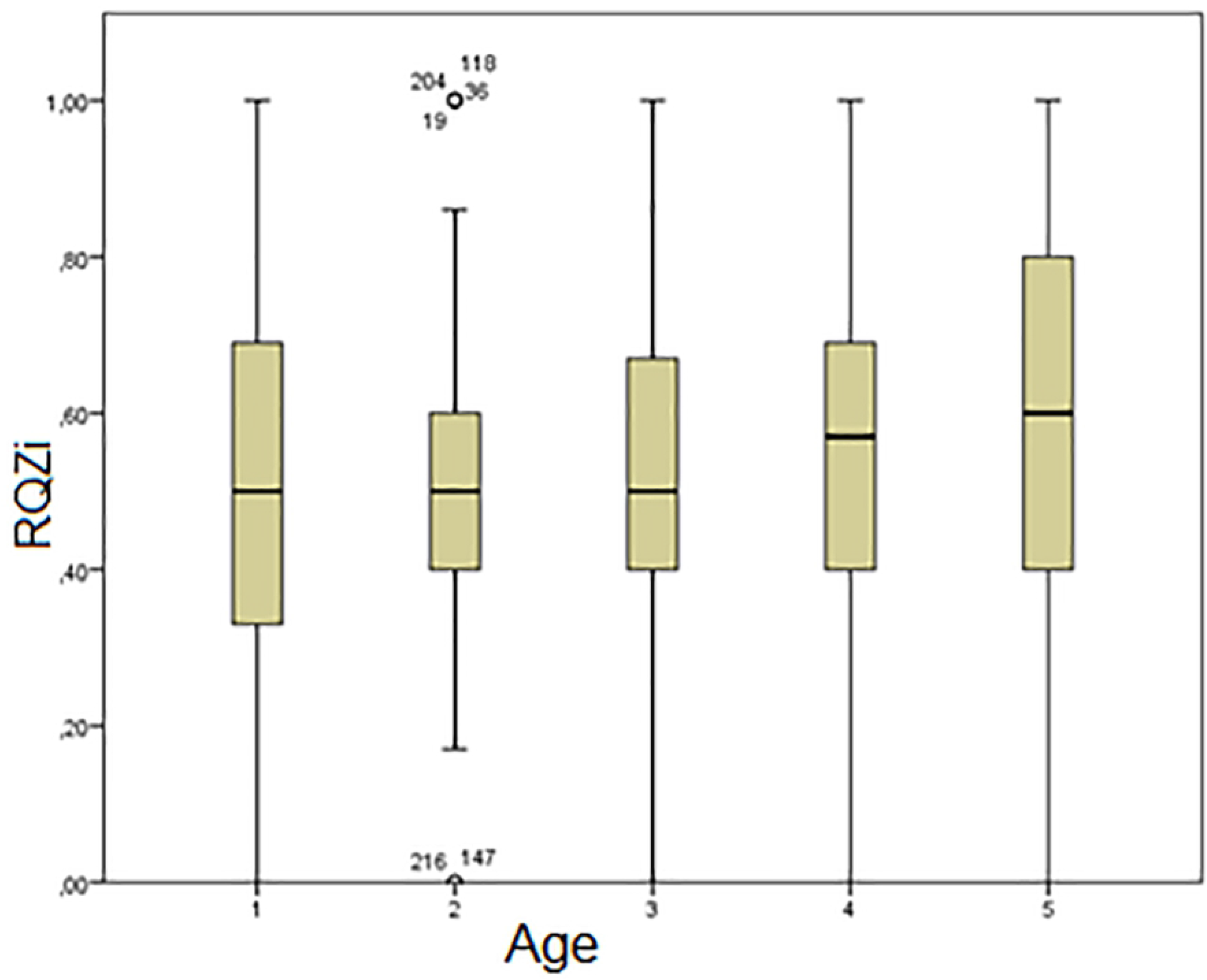
Relationship between age and knowledge index. The ranges were estimated as follows (1) of 30 years or less. (2) 31-39 years old. (3) 40-49 years old. (4) 50 to 59 years old. (5) 60 years or older.

In the 31-39 age group, outliers were reported (Fig 5), ranging from zero, not recognizing the species present in their territory (147 and 216), and other values equal to or close to one (19, 36, 118, and 204), that is, they recognized all or a large number of species, demonstrating a high level of heterogeneity in terms of the RQZi.

### Gender

There is a statistically significant relationship, Chi=6.50 (0.090), between gender and categories of use (Table 4), and with the number of parts used, Chi=8.46 (0.037), where on average women use more plant parts than men (3.4 and 3.2 respectively). Women’s predominant uses are medicinal (69.1%) and ornamental (54.6%), since they are more involved in household activities, while men’s predominant uses are agro-ecological (52.2%) and agricultural (75%) since they are mostly involved in farming and trade.

Our results show no significant differences between the female and male knowledge indexes. Women obtained an average of 0.56 (±0.257), which implies a variation coefficient of 45.5%, and men obtained an average of 0.54 (±0.246), with a variation coefficient of 46.1%. However, it can be seen each municipality has its tendency, in Cucaita, men are the most knowledgeable, while in Ráquira and Villa de Leyva it is women (Fig 6a).

**Fig 6.**
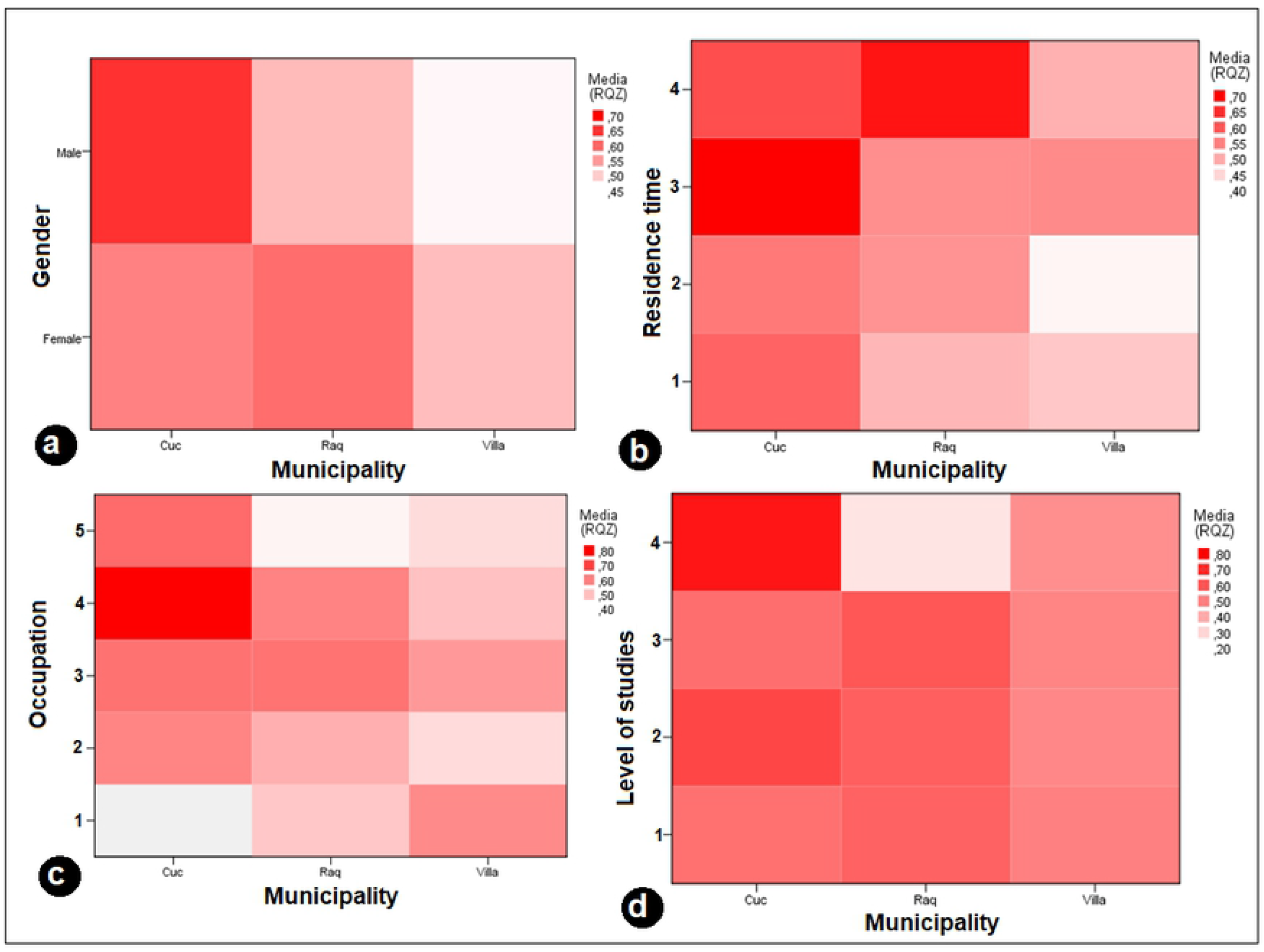
Relationship of socioeconomic factors and the knowledge wealth index for each municipality. a. Gender. b. Time of residence: 20 years or less (1), 21 to 30 years (2), 31 to 50 years (3), 51 years, or more (4). c. Occupation: artisan (1), farmer (2), household trade (3), merchant (4), others (5). d. Level of education organized in four categories: no study (1), primary (2), secondary (3), technical or university (4).

[36] found that men and women present the same knowledge, but differ in the type of categories reported, as men are more dedicated to cultivation and forage, and women are related to food preparation and personal hygiene.

### Level of studies

There is no statistically significant relationship between the level of study with the wealth index knowledge index, number of plants and parts used, or categories of use. However, it should be pointed out that while in some municipalities such as Cucaita there is a tendency that the higher the level of study, the higher the level of knowledge, in Ráquira this behavior is inverse, which indicates the higher level of study, the lower level of knowledge of plants.

In Cucaita, informants with some degree of technical or university level were those who obtained a higher index of knowledge, RQZi between 0.7 to 0.8, concerning people who had a lower degree of schooling, RQZi between 0.4 to 0.6; people with high levels of training, have been able to move to the city of Tunja (the capital of Boyacá) to complete their studies, without the need to migrate completely, due to its proximity (18 km, approximately 30 minutes by car), which makes them not lose their relationship with their local rural environment.

In Villa de Leyva, regardless of the study level, the RQZi remains constant, 0.4 and 0.5. In Ráquira, the RQZi is higher in informants who have some degree of secondary study, RQZi between 0.5 and 0.6, than in interviewees who have reached a technical or university level, RQZi=0.2 (Fig 6d).

In Argentina, people with higher education who know more about medicinal plants, since they periodically carry out training in traditional medicine; also, the knowers, who do not have a level of study, have acquired this knowledge, since they have developed in their rural environment, allowing the transmission of traditional knowledge [73]. [74] have shown that traditional knowledge can often have a negative relationship with the level of schooling since children or adolescents migrate from their natural and cultural environment to receive a formal education, which prevents them from participating in local activities related to the transmission of traditional knowledge by elders.

### Occupation

34.7% of the total number of interviewees are farmers, 33.2% are dedicated to the household trade, 5% are artisans, 5% are merchants, 1% are independent and the remaining percentage is dedicated to other trades, such as construction, electrical network, mining, spinning wool, welding, driving, tour guide, and security guard.

Most farmers were found in Cucaita (44/91) followed by Villa de Leyva (31/91); in Ráquira and Villa de Leyva they were mostly traders (5/13 and 6/13 respectively) and artisans (7/13 and 6/13) respectively). These tasks coincide with the type of economy which is managed in each of these municipalities, Cucaita being mainly agricultural, and Ráquira and Villa de Leyva touristic ones, and similar commerce; however, this type of economic activity affects the human-nature relationship, since people are dedicated to new sources of economic income, generating increasing urbanization and loss of local biodiversity [51].

It was expected in the three municipalities, that farmers would present a higher level of knowledge because of their constant relationship with their territory, however, it was the people involved in commerce, home trades, and handicrafts, the ones who obtained a greater knowledge score for these species (Fig 6c); Cucaita’s commercial activities are mainly focused on the trade of cultivated products, and there is evidence of *O. ficus-indica* cultivation, people relate these species to commercial activity and recognize them easily; people in the home relate mainly to ornamental and food species, and some artisans express they are in search of natural materials for manufacturing their products. They also comment factories are located in rural areas, reasons that keep them in frequent contact with their local biodiversity.

### Residence time living in the area

There is a statistically significant relationship between the number of plants used and the wealth of knowledge index. Informants who have resided in the area for most of their lives, from 31 to 50 years or even more, have the greatest knowledge of cactus species (Fig 6b). In the semi-arid areas of Brazil, [67] also found that those who have resided in the area for the longest time are the ones who know the native food species the most and argue this relationship is because of time interaction between the inhabitants and the resource.

Factors which statistically influence the wealth index of knowledge were age and time of residence in the area; that is to say that people with ages between 40 and 60 or more, and who have resided in the area for a longer time, are the ones who recognize the species of the cactus present in their municipality the most.

However, [68] consider that socioeconomic factors such as education, gender, and occupation are not sufficient to explain the local importance of native plant species in a socioecological system; these authors express that other historical, cultural, ecological, and psychological factors should be taken into account, which can interfere in the socioecological system and would better explain the local importance of the species.

The management and knowledge of cacti in each municipality varied despite being geographically close (approximately 170 km2 was the area sampled) and presenting similar plant formations such as xerophytic and sub xerophytic scrub; in general, Cucaita presented the highest averages of knowledge wealth index according to factors such as gender, occupation, time of residence in the area and educational level, unlike Ráquira and Villa de Leyva, in the latter the lower values of the index were observed (Fig 6).

Cucaita is a municipality with an agricultural tradition, relatively small in extension (43. 6 km2) concerning the other two municipalities, and where the rural population interviewed had some kind of interpersonal relationship and knew each other; Ráquira and Villa de Leyva are larger (233 km2 and 128 km2 respectively), and most of the interviewees did not know each other; This may explain why the smaller the municipality or the urban area, the closer the social network and the greater the interaction between the inhabitants, which facilitates the transfer of knowledge; unlike the larger municipalities, where homes are more separate and there are usually no close family relationships.

Urbanization increasing also brings a work diversification among inhabitants, especially in the municipalities of Ráquira and Villa de Leyva that present a great touristic activity, emphasized in commerce within the urban area, like craft sale, gastronomy, and plans outdoors but in modified environments for the enjoyment of visitors; the previous thing could affect knowledge of its local biodiversity since its economic income doesn’t depend directly on it. It is worth noting that in Villa de Leyva, foreign people have immigrated, changing cultural aspects of this territory.

It is important to keep in mind that the extension of semi-arid zones varies in each municipality and these also have other humid ecosystems where the traditional crops characteristic of the department are grown, such as onions, potatoes, peas, corn, barley, fruits, and tomatoes; the above according to the development plans, to meet the demand and national and international interests [13–15], undervaluing the semi-arid territories and generating people who live there to move daily to work as laborers in these productive sectors.

Following the proposal of [66], conservation strategies for these species should be specific to each municipality due to the influence of the factors mentioned above. Recent studies in Colombia highlight the use of cultivated and wild plants in peasant communities because they play a fundamental role in self-consumption and local agro-biodiversity, which sustains family nutrition. They also highlight the importance of valuing and integrating peasant’s knowledge into ecosystem biodiversity management and conservation plans [75,76].

It should be noted that at the global level and in different studies there has been evidence of a gradual decline in traditional and local knowledge, due not only to the influence of socioeconomic factors of the inhabitants but also to the interaction with other situations such as economic globalization, market interests, lack of interest for new generations and the migration of the rural population to large cities [75,77].

## Conclusions

Of the nine species found, *O. ficus-indica* stands out for having the highest use value and relative importance. *O. soederstromiana* is the second most used species and could be constituted as a potential economic species in the region; the remaining seven species are occasionally valued, for informal sale or fruit consumption at home.

The most-reported use category was food, especially the fruits of *O. ficus-indica* and *O. soederstromiana*, secondly ornamental, of *M. columbiana, M. andinus*, and *M. curvispinus*; and thirdly medicinal use, the stems and flowers of *O. ficus-indica* and *O. soederstromiana*.

Factors such as age and time of residence in the area, affect the local knowledge of cacti, which has been lost in the younger generations and has been maintained by people who have lived much or all of their lives in these territories. Other factors such as gender, educational level, and occupation were not determining factors in the general ethnobotanical knowledge of the area of study, but they do influence in some way the use of these plants in each municipality; in Cucaita, a traditionally agricultural municipality, the highest values of knowledge were registered; followed by Ráquira and Villa de Leyva, where the high tourist and commercial activity has an impact.

The extraction of wild individuals from their habitat of *Mammillaria columbiana, Melocactus andinus*, and *M. curvispinus*, for ornamental purposes, has them seriously threatened. Also, the populations of *C. tunicata* have been reduced because the inhabitants consider it dangerous. Therefore, this study highlights the importance of making a conservation plan for these species, which allows its sustainable use.

Most of the interviewees recognize less than half of the species since they are dedicated to the cultivation of other commercial species (especially *Allium cepa, Allium fistulosum*, and *Solanum tuberosum*), established in humid zones; this emphasizes the need to make the inhabitants aware that it is more profitable in economic terms to use both ecosystems than just using one of it.

## Acknowledgements

To all the farmers from the municipalities of Cucaita, Ráquira, and Villa de Leyva who responded to the interviews. To Yamith Vega, Andrés Espino, and Carlos Alberto Saba, for their accompaniment in the explorations.

This research is part of the project with institutional code SGI 2707 in the Research Management System, evaluated and approved by the Vicerectoría de Investigación y Extensión-VIE of the Universidad Pedagógica y Tecnológica de Colombia.

## Supporting information

**S1 Appendix**. Photographic catalogs of the species shown to informants for each municipality.

**S2 Appendix**. Semi-structured interview.

**S3 Appendix**. Relationships between ethnobotanical data and socioeconomic factors.

